# Anticancer drug doxorubicin spontaneously reacts with GTP and dGTP

**DOI:** 10.1101/2022.10.07.511347

**Authors:** German Mejia, Linjia Su, Popular Pandey, Kevin Jeanne Dit Fouque, Anthony McGoron, Francisco Fernandez-Lima, Jin He, Alex Mebel, Fenfei Leng

**Affiliations:** Biomolecular Sciences Institute, Florida International University, Miami, FL 33199, United States; Department of Chemistry and Biochemistry, Florida International University, Miami, FL 33199, United States; Department of Physics, Florida International University, Miami, FL 33199, United States; Department of Biomedical Engineering, Florida International University, Miami, FL 33199

**Keywords:** anticancer drug, doxorubicin, GTP, dGTP, the doxorubicin-GTP and -dGTP conjugates, ATP

## Abstract

Here we reported a spontaneous reaction between anticancer drug doxorubicin and GTP or dGTP. Incubation of doxorubicin with GTP or dGTP at 37 °C or above yields a covalent product: the doxorubicin-GTP or -dGTP conjugate where a covalent bond is formed between the C14 position of doxorubicin and the 2-amino group of guanine. Density functional theory calculations show the feasibility of this spontaneous reaction. Fluorescence imaging studies demonstrate that the doxorubicin-GTP and -dGTP conjugates cannot enter nuclei although they rapidly accumulate in human SK-OV-3 and NCI/ADR-RES cells. Consequently, the doxorubicin-GTP and -dGTP conjugates are less cytotoxic than doxorubicin. We also demonstrate that doxorubicin binds to ATP, GTP, and other nucleotides with a dissociation constant (K_d_) in the sub-millimolar range. Since human cells contain millimolar levels of ATP and GTP, these results suggest that doxorubicin may target ATP and GTP, energy molecules that support essential processes in living organisms.

---

Doxorubicin is a clinically important anticancer drug widely used in cancer chemotherapy^1-5^. It is approved to treat acute lymphoblastic leukemia (ALL), acute myeloid leukemia (AML), Hodgkin’s lymphoma, breast cancer, and certain metastasized cancer types including stomach cancer, small cell lung cancer, non-small cell lung cancer, and soft tissue and bone sarcomas (https://www.cancer.gov/about-cancer/treatment/drugs/doxorubicinhydrochloride). Doxorubicin is a DNA intercalator with high binding affinity to DNA^6^ and a topoisomerase II poison that stabilizes the gyrase-DNA cleavage-complexes and causes DNA double-stranded breaks^7-9^. It is believed that topoisomerase II poisoning and DNA intercalation are the main mechanism of action (MoA) for its anticancer activities^9-12^. Doxorubicin is a quinone (Fig. 1a) and can be induced to produce reactive free radicals under certain conditions especially in mitochondria^13, 14^. This property is considered to be one of the primary mechanisms responsible for doxorubicin-induced cardiotoxicity^13, 15-18^, a major side effect that limits its therapeutic potential as an anti-cancer drug.

**Figure 1.**
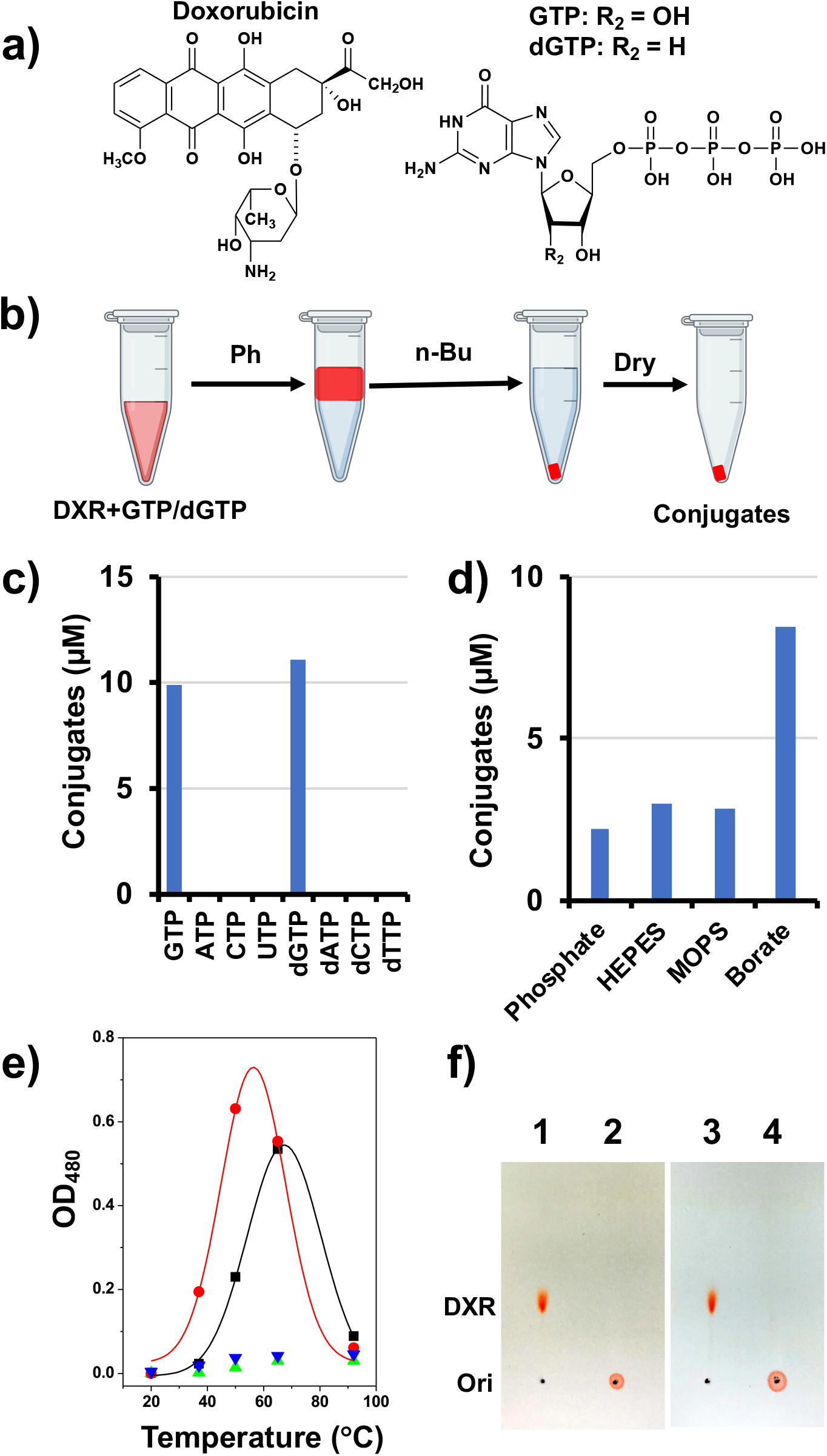
Doxorubicin spontaneously reacts with GTP and dGTP to produce the doxorubicin-GTP and -dGTP conjugates. For a typical reaction, 100 μM doxorubicin and 1 mM nucleotides were used. **a)** Chemical structures of doxorubicin and GTP/dGTP. **b)** An experimental procedure to synthesize doxorubicin-GTP and –dGTP conjugates. **c)** Only GTP and dGTP reacted with doxorubicin spontaneously in 10 mM borate buffer, pH 8.2 at 37 °C. **d)** Doxorubicin was conjugated to GTP in 50 mM sodium Phosphate buffer, pH 7.4, 50 mM HEPES-KOH buffer, pH 7.5, 10 mM MOPS buffer pH 8.0, and 10 mM Borate buffer, pH 8.2. **e)** The spontaneous reaction of doxorubicin and GTP at 20, 37 52, 65, and 90 °C for 2 (black squares) and 12 (red circles) hours. Up and down triangles represent doxorubicin in the absence of GTP and dGTP. **f)** Thin layer chromatography (TLC) on silica gel 60 for doxorubicin (lanes 1 and 3) and the GTP-doxorubicin conjugate (lanes 2 and 4). After the samples were spotted, the TLC plates were developed with a solvent system of chloroform : methanol : acetic acid = 16 : 4 : 1 (v/v). Samples in lanes 3 and 4 were incubated at 65 °C for 2 hours.

Although doxorubicin has been studied extensively^4, 19-22^, the chemical properties and biological effects of this anticancer compound are still not fully understood. For instance, previous studies showed that doxorubicin binds to ATP and other nucleotides *in vitro*^23, 24^. However, a systematic investigation how doxorubicin interacts with these energy-rich molecules has not been conducted yet. The reactivities of doxorubicin toward these nucleotides have also not been examined. Since ATP and GTP are two main energy molecules that drive most vital processes in live cells, binding and reaction with them should significantly compromise their availability and functions, and is expected to have grave consequences. Here, we report a new spontaneous chemical reaction between doxorubicin and GTP or dGTP: simple incubation of doxorubicin with GTP or dGTP at 37 °C yields two covalent products, the doxorubicin-GTP and -dGTP conjugates. Our results show that the covalent bond forms between the C14 of doxorubicin and the 2-amino group of guanine. We also demonstrate that doxorubicin binds to GTP, dGTP, ATP, and dATP with a dissociation constant (K_d_) in the sub-millimolar range, suggesting that doxorubicin may target these energy molecules inside cells.

## Results and Discussion

Fig. 1b shows a simple experimental procedure in which mixing of doxorubicin with GTP or dGTP spontaneously produces two covalent products, the doxorubicin-GTP and -dGTP conjugates. Initially, we used a borate buffer (10 mM sodium borate, pH 8.2) for this reaction and tested reactivities between doxorubicin and one of the 8 nucleotides, GTP, ATP, CTP, UTP, dGTP, dATP, dCTP, and dTTP. Fig. 1c shows our results. Apparently, only GTP and dGTP spontaneously reacted with doxorubicin and the other 6 nucleotides did not react with doxorubicin at all. We also tested different buffers for this spontaneous reaction and found that doxorubicin reacted with GTP or dGTP in all four tested buffers (Fig. 1d). The best buffer was borate buffer which yielded more products. Since the borate buffer has a higher pH, it is possible that the basic condition favored the spontaneous reaction. More experiments are needed to confirm this hypothesis. We also performed this experiment in different temperatures and found that temperature greatly affected this spontaneous reaction (Fig. 1e). Doxorubicin did not react with GTP or dGTP at 20 and 90 °C. Significantly more doxorubicin-GTP and -dGTP were produced at ∼60 °C. Nevertheless, the fact that doxorubicin spontaneously reacts with GTP and dGTP at 37 °C suggests that this anticancer drug reacts with GTP and dGTP under physiological conditions. It is likely that doxorubicin reacts with GTP and dGTP in cancer cells of patients. Since doxorubicin-GTP and -dGTP conjugates contain four negative charges, they did not migrate from their origins in the thin layer chromatography (TLC) plates under the experimental conditions used in this study (Fig. 1f, lanes 2 and 4). We also incubated the doxorubicin-GTP conjugate at 65 °C for 2 hours and found that doxorubicin-GTP conjugate was stable and did not decompose into doxorubicin and GTP (Fig. 1f, lane 4).

Next, we generated sufficient amounts of the doxorubicin-GTP conjugate and purified the product using a C18 solid phase extraction (SPE) column. A Fourier-transform ion cyclotron resonance (FTICR) mass spectrometry was performed to confirm the chemical structure/identity of the doxorubicin-GTP conjugate. The experimental results (Fig. 2a) are consistent with theoretical prediction, indicating that the linkage is between the C14 of doxorubicin and the 2-amino group of guanine (Fig. 1a and Fig. 2b). Daunorubicin cannot react with GTP or dGTP further confirming that the spontaneous reaction is between the C14 of doxorubicin and the 2-amino group of guanine. Fig. 2c and d show the visible absorbance and fluorescence emission spectra of doxorubicin and the doxorubicin-GTP conjugate. The conjugation of doxorubicin to GTP results in a red shift of the absorbance maximum from 480 to 505 nm, similar to doxorubicin binding to DNA ((56); Fig. 2c). The fluorescence intensity was also significantly decreased (Fig. 2d).

**Figure 2.**
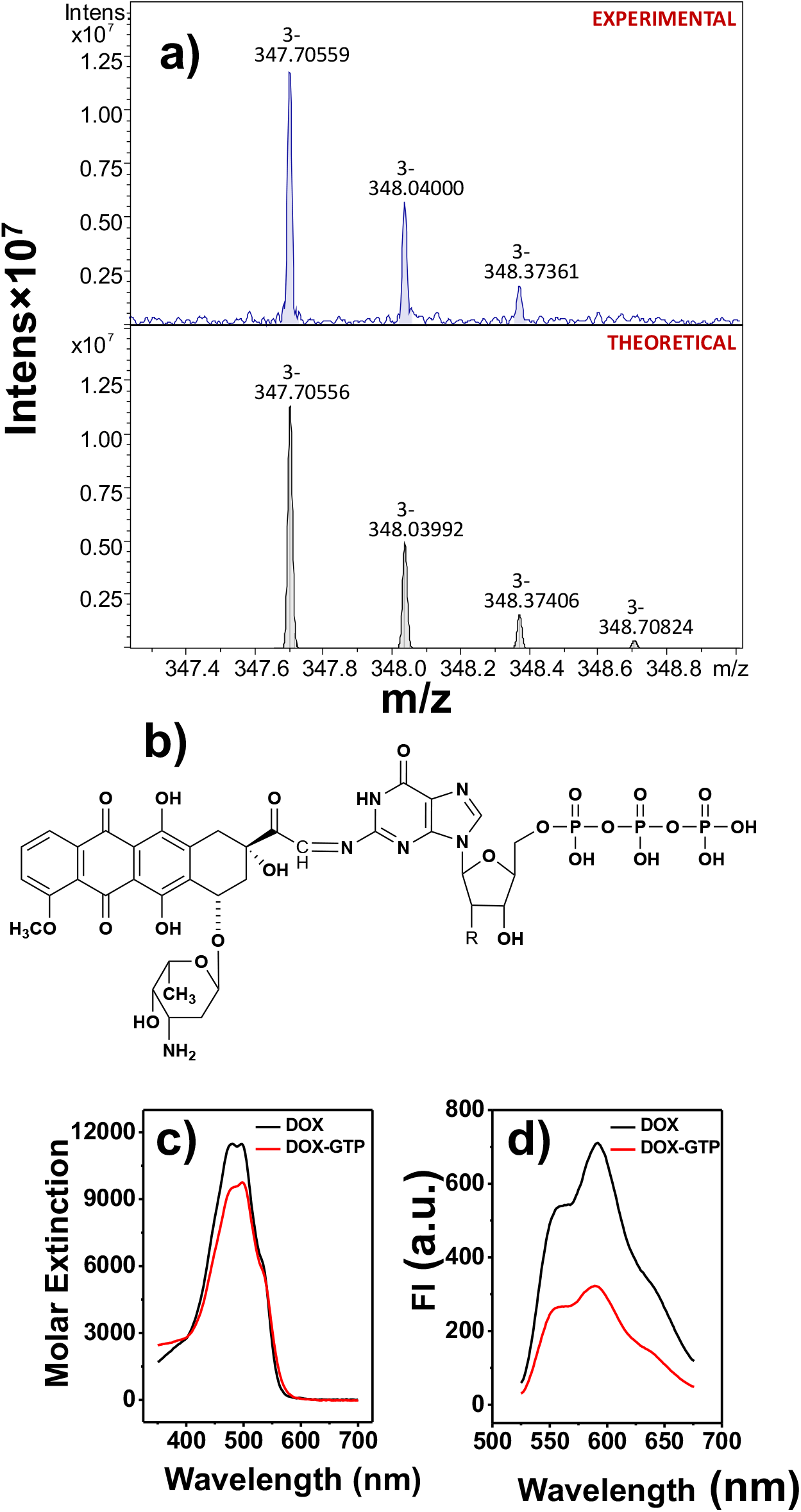
**a)** The experimental (upper panel) and theoretical (lower panel) FTICR mass spectrometry spectra of the doxorubicin-GTP-2 conjugate. **b)** Deduced chemical structure of doxorubicin-GTP according to the mass spectrometry results. **c)** Visible absorbance spectra of doxorubicin (black line) and the doxorubicin-GTP-2 conjugate (red line) in 1×BPE buffer. **d)** The fluorescence emission spectra of doxorubicin (black line) and the doxorubicin-GTP-2 conjugate (red line) in 1×BPE buffer with the excitation wavelength of 480 nm.

A requirement for the spontaneous reaction between doxorubicin and GTP is that doxorubicin must bind to GTP under our experimental conditions. In this study, visible absorbance and fluorescence titration experiments were used to investigate how doxorubicin binds to GTP. Fig. 3 shows our results. Titrating GTP into a fixed concentration of doxorubicin results in a decrease of absorbance around 480 nm and an increase of absorbance around 600 nm. The existence of an isosbestic point indicates a simple, two-state binding reaction (Fig. 3a). Fitting the binding data to a simple binding model yields a binding constant of 3.52±0.51×10^4^ M^-1^ and a 1-to-1 binding stoichiometry. The 1-to-1 binding stoichiometry is confirmed by the Job plot of doxorubicin binding to GTP. Similar results were also obtained with fluorescence titration experiments. Titrating GTP into a fixed concentration of doxorubicin dramatically quenched the fluorescence of doxorubicin (Fig. 3d). Interestingly, fitting these results to a simple binding model yields a binding constant of 0.88±0.22×10^3^ M^-1^ and a 1-to-1 binding stoichiometry (Fig. S1a). The binding constant is significantly lower than the one obtained from the absorbance titration experiments. The 1-to-1 binding ratio was confirmed by the fluorescence Job plot of doxorubicin binding to GTP (Fig. S1b). Further studies are needed to determine the discrepancy between the absorbance and fluorescence titration experiments. Nevertheless, absorbance and fluorescence titration experiments show that doxorubicin binds to GTP with a dissociation constant (K_d_) in the sub-millimolar range in aqueous buffer solutions. In this study, we also conducted absorbance and fluorescence titration experiments for several other nucleotides including dGTP, GMP, dGMP, ATP, dATP, and etc. Our results are summarized in Fig. S2 and Table S1. Doxorubicin binds all nucleotides with 1-to-1 binding ratio and similar binding affinities (Table S1).

**Figure 3.**
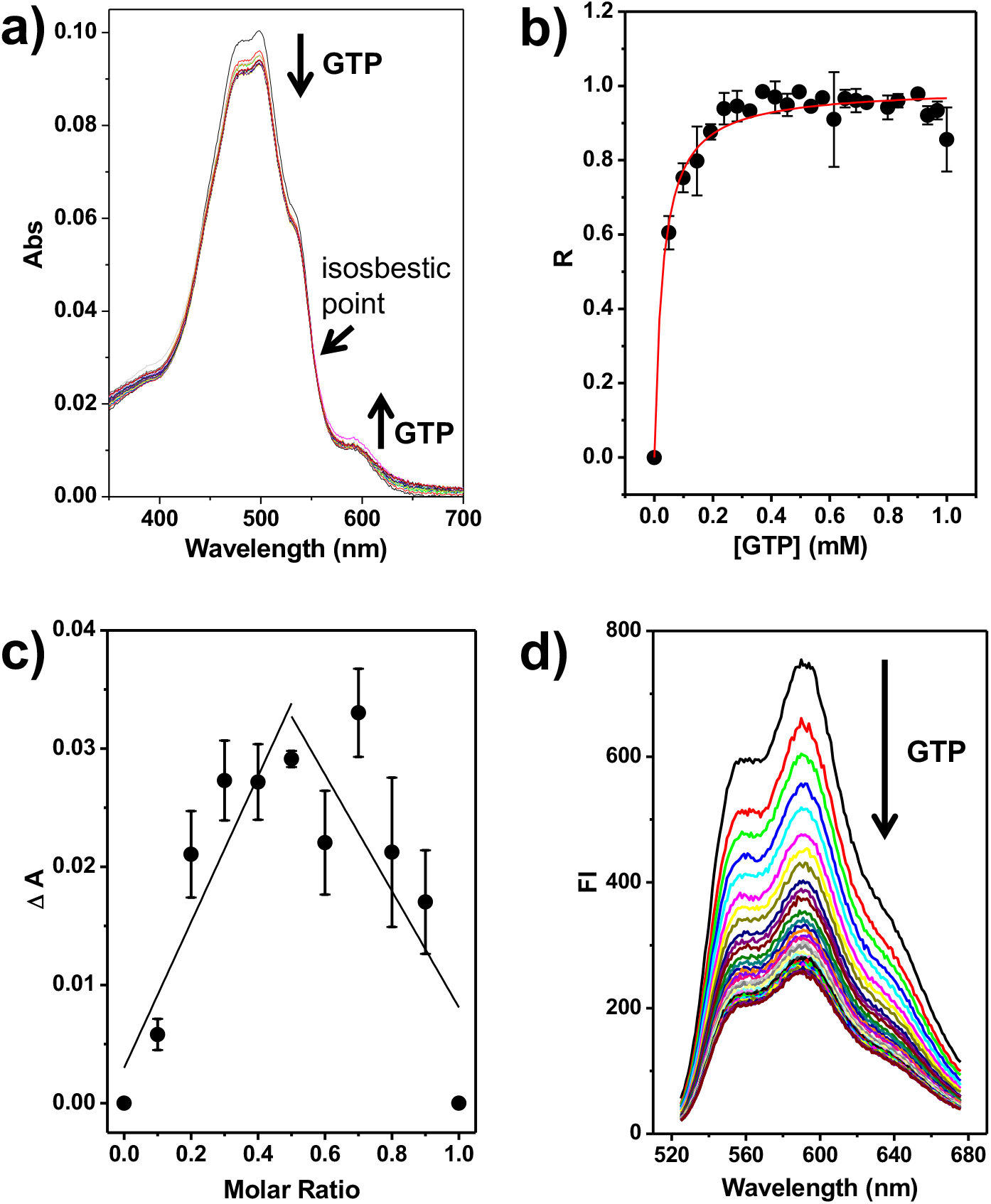
Binding of doxorubicin to GTP in 1×BPE buffer. **a)** Absorbance titration experiments were performed as described under Materials and Methods. **b)** The binding curve is obtained by nonlinear-least-squares fitting of equation (1) as described under Materials and Methods. The binding constant of doxorubicin binding to GTP was determined to be 3.6 ±0.8 ×10^4^ M^-1^. **c)** The Job plot for doxorubicin binding to GTP in in 10 mM sodium borate buffer pH 8.2. **d)** The fluorescence titration of doxorubicin to GTP. Error bars represent standard deviations from three independent experiments.

The noncovalent binding of doxorubicin to GTP and dGTP was also confirmed by mass spectrometry studies. In fact, the nESI-MS analysis of the DXR·GTP (1066.2 Da, Fig. 4a) and DXR·dGTP (1050.2 Da, Fig. 4b) complexes in an equimolar ratio both resulted in a 1:1 binding stoichiometry with a single charge state isotopic pattern under native-like solution conditions. In addition, similar native nESI-MS experiments using different nucleotides, consisting of DXR·GMP (906.2 Da, Fig. 4c), DXR·ATP (1050.2 Da, Fig. 4d) and DXR·AMP (890.3 Da, Fig. 4e), were also found to bind to doxorubicin in a 1:1 binding stoichiometry. The present gas-phase mass spectrometry results were consistent with the solution-phase observations. In this MS study, ion mobility experiments were also conducted for each observed DOX complexes. The TIMS spectra exhibited a characteristic collision cross section (CCS) of ∼285 Å^2^ for DXR·GTP, DXR·dGTP and DXR·ATP complexes, indicating that the GTP, dGTP and ATP nucleotides adopt similar gas-phase conformation when binding to doxorubicin (Fig. 4, right panel). The same observations were obtained for the DXR·GMP and DXR·AMP complexes, for which a characteristic CCS of ∼265 Å^2^ was obtained for the DXR·GMP and DXR·AMP complexes.

**Figure 4.**
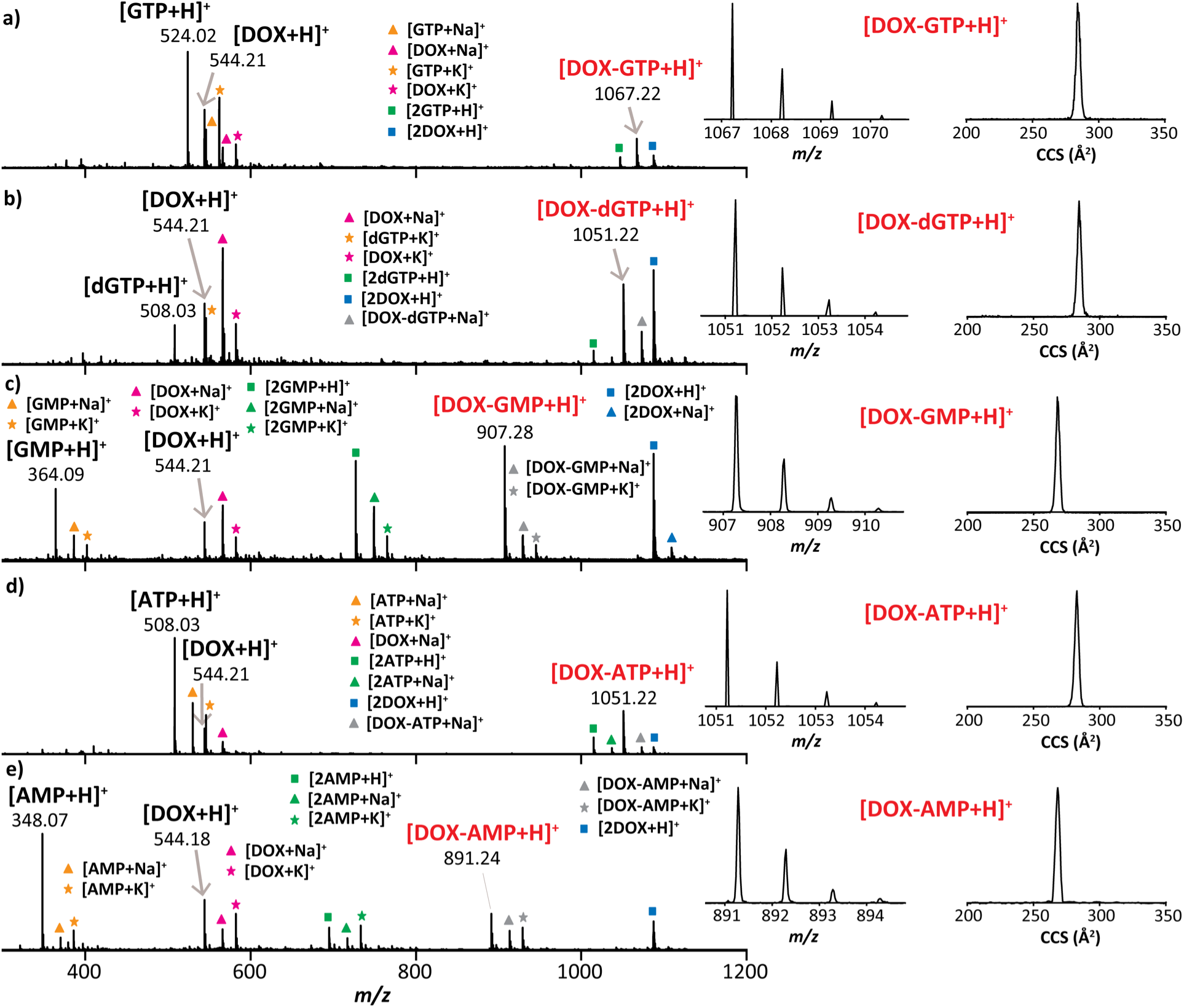
Mass spectrometry analysis showing the native nESI-generated mass spectra, singly charged isotopic patterns and native ion mobility distributions for **a)** DXR:GTP, **b)** DXR:dGTP, **c)** DXR:GMP, **d)** DXR:ATP and **e)** DXR:AMP complexes.

The reaction between doxorubicin and GTP was explored theoretically via density functional theory (DFT) calculations of thermodynamic functions for the formation of a van der Waals complex between the two molecules and a chemically bound product through the formation of a new C-N bond between the amino nitrogen in GTP and the carbon of the CH_2_OH group in doxorubicin with a release of a water molecule. The calculations show that doxorubicin and GTP readily form a van der Waals complex (Fig. 5) with the binding enthalpy and Gibbs free energy at 298.15 K of 16.4 and 1.8 kcal/mol, respectively. The binding is mostly provided by hydrogen bond formation between hydroxyl groups in GTP and doxorubicin and between the amino group of GTP and hydroxyl and oxo groups in doxorubicin, as illustrated in Fig. 5. Further, a covalent C-N bond formation between the amino nitrogen in GTP and the carbon of the CH_2_OH group in doxorubicin with a release of a water molecule results in the chemically bound product. The binding is also slightly enhanced by hydrogen bonding between an OH group in GTP and an oxo group in doxorubicin. The reaction of the chemical product formation along with H_2_O is calculated to be slightly endoergic, by 2.9 and 5.2 kcal/mol in terms of enthalpy and Gibbs free energy, respectively. This indicates that the reaction is nearly thermoneutral within the accuracy of the DFT method used and hence likely feasible even at room temperature and can be further enhanced with a temperature increase. However, further oxidation of the chemical product with a formation of the double C=N and a release of H_2_ is thermodynamically unfavorable, as it leads to a raise of enthalpy and Gibbs free energy to 34.6 and 30.4 kcal/mol, respectively, as compared to the initial reactants. Nevertheless, the alcohol at C13 can be oxidized by O_2_ in aqueous solution to form the double C=N and yield the final product^25, 26^. Thus, the calculation results attest to the thermodynamic feasibility both for the formation of the van der Waals complex and chemically bound product with a single C-N bond and provide structural information for these species.

**Figure 5.**
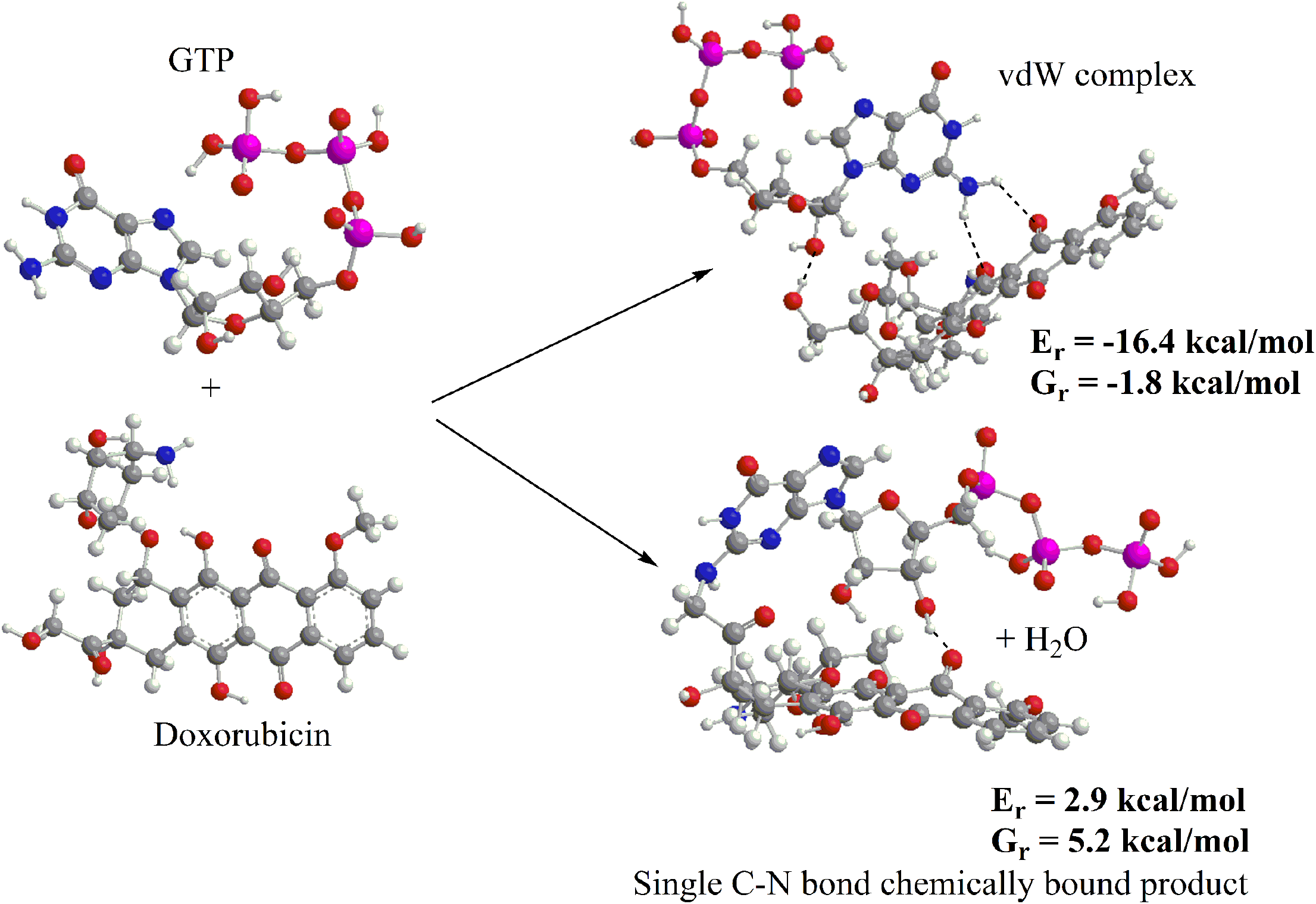
DFT calculations. Geometries of GTP, doxorubicin, their van der Waals complex, and a covalently bond complex optimized at the B3LYP/6-31G(d) level of theory. Reaction energies for the formation of the complexes and the corresponding Gibbs free energies at 298.15 K are also shown.

MTT cell viability assays were used to evaluate the *in vitro* cytotoxicity of doxorubicin-GTP and -dGTP conjugates against two human ovarian cell lines: the doxorubicin-sensitive SK-OV-3 and the doxorubicin-resistant NCI/ADR-RES cell lines^27, 28^. To our surprise, doxorubicin-GTP and - dGTP conjugates are much less cytotoxic to both cell lines than the parent compound, doxorubicin (Fig. 6a). The IC_50_ values of doxorubicin against SK-OV-3 and NCI/ADR-RES cells were determined to be 5.2 ±2.6 ×10^−7^ and 5.8 ±0.8 ×10^−5^ M, respectively, which is consistent with previously published results^27, 29, 30^. In contrast, more than 80% of cells are viable when 10 or 200 μM of doxorubicin-GTP and -dGTP are used in the MTT assays for doxorubicin-sensitive SK-OV-3 and doxorubicin-resistant NCI/ADR-RES cell lines, respectively. We also used fluorescence microscopic imaging to examine whether the doxorubicin-GTP and -dGTP conjugates enter/accumulate in these cancer cells and to identify its cellular distributions/locations. Fig. 6b shows our result. As expected, doxorubicin entered SK-OV-3 cells and accumulated in nuclei rapidly (Fig. 6B). Much less doxorubicin entered/accumulated in NCI/ADR-RES nuclei (Fig. 6b), which is consistent with the published results^29, 30^. In contrast, although both doxorubicin-GTP and -dGTP conjugates enter and accumulate in SK-OV-3 and NCI/ADR-RES cells rapidly, these two new compounds only stay in cytosol and cannot enter cell nuclei. This can explain why cells are viable after the incubation with doxorubicin-GTP and -dGTP conjugates. Further studies are required to determine other biological effects and the long-term cytotoxicity of doxorubicin-GTP and -dGTP conjugates in the future.

**Figure 6.**
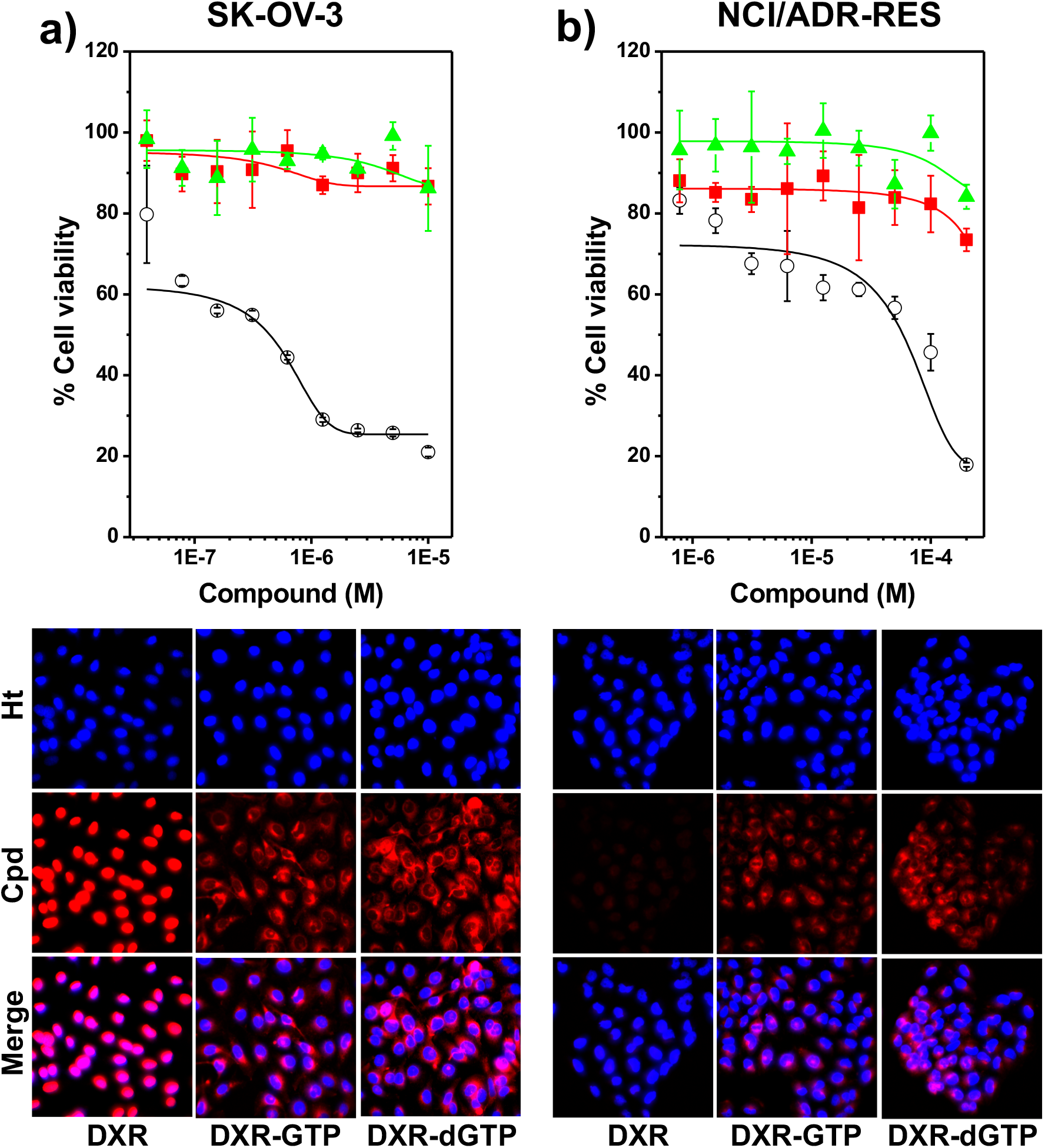
Cytotoxicity of doxorubicin, doxorubicin-GTP or -dGTP against human ovarian SK-OV-3 and NCI/ADR-RES cell lines. Cells were treated with indicated concentrations of doxorubicin, doxorubicin-GTP or -dGTP conjugate for 72 hours. Cytotoxicity was then evaluated by MTT assay as detailed in Materials and Methods. **a)** Doxorubicin-sensitive SK-OV-3 cells. IC_50_ was determined to be 5.2±2.6×10^−7^ M for doxorubicin (black circles). The doxorubicin-GTP (red squares) and –dGTP (green triangles) conjugates did not significantly inhibit cell growth. **b)** Doxorubicin-resistant NCI/ADR-RES cells. IC_50_ was determined to be 5.8±0.8×10^−5^ M for doxorubicin (black circles). The doxorubicin-GTP (red squares) and –dGTP 3(green triangles) conjugates did not significantly inhibit cell growth. All values are expressed as the mean ±SD for triplicated samples. Fluorescence microscopy images of SK-OV-3 **(a, left)** and NCI/ADR-RES **(b, right)** cells after incubation with 5 μM of doxorubicin (DXR), the doxorubicin-GTP conjugate (DXR-GTP), the doxorubicin-dGTP conjugate (DXR-dGTP), respectively for 4 h. The images were taken under 20×. Ht and Cpd represent hoechst33258 and compound, respectively.

The spontaneous reaction between doxorubicin and GTP/dGTP, and the tight binding of doxorubicin to GTP/ATP suggest that doxorubicin, in addition to intercalating into DNA base pairs and poisoning DNA topoisomerase II, also targets ATP/dATP and GTP/dGTP inside cells. Doxorubicin has two subcellular locations: nuclei and mitochondria^13, 18, 31-34^. Previous studies showed that the average concentration of GTP in human cells is 468±224 μM^35^. Mitochondria also contain millimolar levels of GTP, GDP, and GMP^36^. Since normal human body temperature is 37 °C, it is feasible for doxorubicin to spontaneously react with GTP and dGTP inside cells and yield the doxorubicin-GTP and -dGTP covalent products when doxorubicin is used for cancer chemotherapy. This covalent reaction should be more favorable in mitochondria since the pH of mitochondrial matrix is higher^37^. The tight binding of doxorubicin to ATP, GTP, and other nucleotides is another concern for the use of doxorubicin as an anticancer drug. In addition to GTP, human cells also contain millimolar levels of ATP, UTP, CTP, and other nucleotides^35, 38^. The high concentrations of intracellular nucleotides provide a condition for doxorubicin binding to these nucleotides in cells, which significantly compromises the utility of ATP and GTP as energy molecules to support their cellular functions. This compromise is a serious problem for cardiomyocytes containing substantial amounts of mitochondria that occupy ∼30% of cardiac cell volume^39^ and produce >95% of ATP responsible for the orchestrated contraction–relaxation cycle of a healthy heart^40^. In other words, the spontaneous reaction between doxorubicin and GTP/dGTP, and the tight binding of doxorubicin to GTP/ATP may significantly contribute to the doxorubicin-induced cardiotoxicity. Further studies are required to investigate this hypothesis.

In summary, a new spontaneous reaction between doxorubicin and GTP or dGTP is discovered. Incubation of doxorubicin with GTP or dGTP at 37 °C and above yields two covalent products: doxorubicin-GTP and -dGTP conjugates. Our results show that the covalent bond forms between the C14 of doxorubicin and the 2-amino group of guanine. DFT calculations show the feasibility of this spontaneous reaction. Fluorescence imaging studies show that doxorubicin-GTP and - dGTP conjugates cannot enter nuclei although they rapidly enter and accumulate in human SK- OV-3 and NCI/ADR-RES cells. Utilizing combined biophysical methods including absorbance and fluorescence titration assays, mass spectrometry, and theoretical calculation, we demonstrate that doxorubicin tightly binds to ATP, GTP, and other nucleotides. These results suggest that doxorubicin may target ATP and GTP *in vivo* and limit their availability as energy molecules. This study raises a safety concern for the use of doxorubicin as an anticancer drug.

## Materials and Methods

### Materials

Doxorubicin was purchased from Waterstone Technology, LLC. A molar extinction coefficient of 11,500 M^-1^ cm^-1^ at 480 nm was used to determine the concentrations of doxorubicin^41^. Adenosine-5’-triphosphate disodium (ATP) and guanosine-5’-triphosphate sodium salt (GTP) were purchased from Amersham Pharmacia Biotech, Inc. The molar extinction coefficients of 15,400 M^-1^ cm^-1^ at 259 nm and 13,700 M^-1^ cm^-1^ at 253 nm, respectively, were used to determine the concentrations of ATP and GTP. Cytidine-5’-triphosphate (CTP) and uridine 5’-triphosphate (UTP) were purchased from New England Biolabs, Inc. 2’-deoxyadenosine-5’-triphosphate (dATP), 2’-deoxyguanosine-5’-triphosphate (dGTP), 2’-deoxycytidine-5’-triphosphate (dCTP), thymidine-5’-triphosphate (dTTP), Gibco™ RPMI 1640 Medium (11-875-093), Corning™ Cellgro™ Cell Culture Phosphate Buffered Saline, Corning™ Penicillin-Streptomycin Solution, Corning™ 0.25% Trypsin and 0.1% EDTA in HBSS without Calcium, and BioLife 35mm Tissue Culture Dish were purchased from Thermo Fisher Scientific, Inc. Sodium borate decahydrate (Na_2_B_4_O_7_·10 H_2_O), 1-butanol, and chloroform (99.9+%, A.C.S. HPLC grade) were purchased from Sigma Aldrich Chemical Co., Inc. Plasmocin™ prophylactic (ant-mpp) was purchased from InvivoGen. HyClone™ standard fetal bovine serum (FBS, SH30088.03) was purchased from GE Healthcare Life Sciences. 96-Well CytoOne®plate, TC-treated (CC7682-7596) was purchased from USA Scientific, Inc. 3-(4,5-dimethylthiazol-2-yl)-2,5-diphenyltetrazolium bromide (MTT) was purchased from Tocris Bioscience. SK-OV-3 human ovarian cancer cells were obtained from American Type Culture Collection (Manassas, VA). NCI/ADR-RES DXR resistant ovarian cancer cells were a gift from Dr. Daping Fan at the University of South Carolina.

### Synthesis and purification of the doxorubicin-GTP and -dGTP conjugates

For a typical conjugation experiment, 100 μM of doxorubicin and 1 mM of GTP or dGTP were incubated at 37 °C in 10 mM sodium borate buffer pH 8.2. After 24 hours of incubation or specified time, reactions were stopped by two times of phenol extraction. The products were precipitated with 10 volumes of 1-butanol, dried using a CENTRIVAP concentrator, and stored at a -80 °C freezer until use. C18 solid phase extraction (SPE) columns may be used to further purify the products.

### Thin Layer Chromatography (TLC)

Doxorubicin and derivatives were spotted on TLC Silica Gel 60 F_254_ and developed with a solvent consisting of chloroform: methanol: acetic acid = 16:4:1 (v/v) at room temperature. After development, the TLC Silica Gels were air dried and photographed under visible and UV light.

### Mass spectrometry

Working solutions of 20 ppm the doxorubicin-GTP conjugate were prepared in 1:1 Optima grade Methanol/Water. The samples were directly infused into the ESI source of a Bruker Solarix FT-ICR, operated at 256 KWord in negative ion mode. Scan acquisition of 50 coadded scans was collected. Mass spectrometry was calibrated utilizing Tuning Mix calibration standard in the range between 112 to 1033 with 5 calibration points obtaining a standard deviation of 0.613. A solvent blank was also analyzed as reference. Solvents, methanol utilized in this study were analytical grade or better and purchased from Fisher Scientific (Pittsburgh, PA). A Tuning Mix calibration standard (G24221A) was obtained from Agilent Technologies (Santa Clara, CA) and used as received. Mass spectra were processed using Bruker Compass Data Analysis version 5.1 (Bruker Daltonik GmbH).

To study the binding of doxorubicin to nucleotides, solutions of nucleotides (e.g., GTP, dGTP, ATP, GMP and AMP) in complex with doxorubicin (1:1 ratio) were analyzed at a concentration of 5 μM in 10 mM NH_4_Ac for native conditions. Low concentration Tuning Mix standard (G1969-85000) was used to calibrate the mass spectrometry and ion mobility spectrometry and obtained from Agilent Technologies (Santa Clara, CA). Mass spectrometry experiments were conducted on a Solarix 7T FT-ICR mass spectrometer (Bruker, Billerica, MA) equipped with an Infinity cell and an ESI source operated in negative ion mode. The high voltage, capillary exit, and skimmer I were set to 1400 V, 120 V, and 25 V respectively. A total of 50 scans (*m/z* range 100-2000) were co-added with a data acquisition size of 256k words.

Ion mobility mass spectrometry experiments were performed on a custom built nESI-TIMS coupled to an Impact q-ToF mass spectrometer (Bruker Daltonics Inc., Billerica, MA). nESI emitters were pulled from quartz capillaries (O.D. = 1.0 mm and I.D. = 0.70 mm) using Sutter Instruments Co. P2000 laser puller (Sutter Instruments, Novato, CA). Solutions were loaded in a pulled-tip capillary, housed in a mounted custom built XYZ stage in front of the MS inlet, and sprayed at 800 V via a tungsten wire inserted inside the nESI emitters. Briefly, the ion mobility separation in a TIMS device is based on holding the ions stationary using an electric field (*E*) against a moving buffer gas^42, 43^. The TIMS unit is controlled by an in-house software in LabView (National Instruments) and synchronized with the MS platform controls. The general fundamentals of TIMS as well as the calibration procedure have been described in the literature^44-47^. TIMS experiments were carried out using nitrogen (N_2_) as buffer gas, at ambient temperature (*T*) with a gas velocity defined by the funnel entrance (*P*_*1*_ = 2.5 mbar) and exit (*P*_*2*_ = 0.9 mbar) pressure differences. An rf voltage of 250 Vpp at 880 kHz was applied to all electrodes. Here, a *V*_*def*_ of 60 V with a *V*_*ramp*_ at −250 to -50 V and base voltage (*V*_*out*_) of 60 V was used for DOX·Nucleotide complexes. Mass and ion mobility spectra were processed using Bruker Compass Data Analysis version 5.1.

### Absorbance titration experiments

Absorbance titration experiments using a Varian Cary Bio-50 UV-Vis spectrophotometer were used to study doxorubicin binding to GTP, ATP, and other nucleotides, and to determine their binding constants. Specifically, the visible absorbance of doxorubicin was measured while increasing concentrations of GTP or other nucleotides were titrated into a 10 μM of doxorubicin solution in 1×BPE (6 mM Na_2_HPO_4_, 2 mM NaH_2_PO_4_, and 1 mM Na_2_EDTA, pH 7.0) or 10 Mm sodium borate buffer pH 8.2 at room temperature. The concentrations of free (C_f_) and bound doxorubicin (C_b_) were calculated using these two equations: C_b_ = (A_0_ – A) / (ε_f_ – ε_b_) and C_f_ = C_T_ – C_b_, where A_0_ and A is the absorbance of doxorubicin in the absence and presence of GTP, respectively. C_T_ is the total doxorubicin concentration. ε_f_ and ε_b_ are extinction coefficients of doxorubicin in the absence and presence of nucleotides, respectively. ε_b_ was determined in the presence of a large excess of one nucleotide with no further change in absorbance at 480 nm. The binding ratio R=C_b_/C_T_. The dissociation constant (K_d_) is calculated by nonlinear-least-squares fitting of the flowing equation:

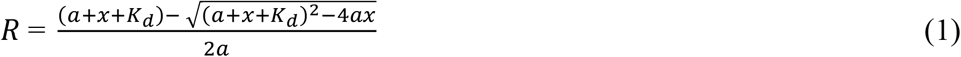

where R, a, and x represented the GTP-binding ratio of doxorubicin, the total doxorubicin concentration, and the total GTP concentration.

### Fluorescence titration experiments

Fluorescence titration experiments using a Varian Cary Eclipse fluorescence spectrophotometer were employed to study how doxorubicin binds to one nucleotide at room temperature with λex= 480 nm and λem= 590 nm. A slit width of 10 nm was used for both excitation and emission. Fluorescence spectra were recorded by titrating one nucleotide into 1 μM of doxorubicin in 1×BPE (6 mM Na_2_HPO_4_, 2 mM NaH_2_PO_4_, and 1 mM Na_2_EDTA, pH 7.0) or 10 mM sodium borate buffer pH 8.2 and used to calculate the concentrations of free (C_f_) and bound doxorubicin (C_b_): C_f_ = C_T_ (I – I_∞_) / (I_0_ – I_∞_) and C_b_ = C_T_ – C_f_, where C_T_ is the total doxorubicin concentration; I_0_ is the fluorescence intensity of free doxorubicin; and I_∞_ is the fluorescence intensity of the doxorubicin in the presence of a large excess of a nucleotide.

### Continuous variation analysis

Binding stoichiometry of doxorubicin to GTP was obtained using the method of continuous analysis^48^. 100 μM of doxorubicin and GTP in the 10 mM sodium borate buffer pH 8.2 were prepared. A series of solution mixtures were made by varying volumes of these two equally concentrated solutions to keep the sum of the concentrations of doxorubicin and GTP constant at 100 μM. The absorbance of doxorubicin in the presence or absence of GTP was measured at 480 nm at room temperature. Similarly, fluorescence intensity at λex = 480 nm with λem = 590 nm was also measured. The difference in absorbance (ΔA) or fluorescence intensity (ΔF) was plotted against the molar fraction of doxorubicin to determine the binding stoichiometry of doxorubicin to GTP.

### Cell culture

SK-OV-3 (doxorubicin-sensitive) and NCI/ADR-RES (doxorubicin-resistant) human ovarian cancer cells were cultured in RPMI 1640 medium supplemented with 10% FBS, 100 IU/mL of penicillin, 100 μg/mL of streptomycin, and 2.5 μg/mL of plasmocin in 25 cm^2^ tissue culture flasks at 37 °C in a humid atmosphere with 5% CO2 and 95% air. The cells were subcultured once they reached 80% confluence and cell density, determined with a hemocytometer prior to each experiment.

### Fluorescence microscopic imaging

SK-OV-3 and NCI/ADR-RES cells (4×10^5^) were seeded in 35 mm tissue culture dishes with 2 mL RPMI 1640 medium supplemented with 10% FBS, 100 IU/mL of penicillin, 100 μg/mL of streptomycin, and 2.5 μg/mL of plasmocin and incubated at 37 °C in a humid atmosphere with 5% CO2 and 95% air for 48 hours. Cells were incubated for additional 4 hours with fresh medium containing 5 μM of doxorubicin, the doxorubicin-GTP conjugate, or the doxorubicin-dGTP conjugate. After the incubation, cells were washed thrice with cold PBS and fixed with 4% paraformaldehyde for 30 min. Next, cells were stained by Hoechst 33258 for 15 min for nuclei and washed thrice with PBS prior to fluorescence imaging. A Home-built inverted fluorescence microscope with an LED light source (X-cite 120 LED Boost, USA) for sample illumination was used to acquire all fluorescence images.

### MTT cell viability assay

Cytotoxicity of doxorubicin, the doxorubicin-GTP conjugate, or the doxorubicin-dGTP conjugate against SK-OV-3 and NCI/ADR-RES cells was evaluated by MTT cell viability assays. Briefly, cells were seeded in 96-well plates (7500 cells/well) in 100 μL of RPMI 1640 medium containing 10% FBS and grew at 37 °C for 24 hours in a humid atmosphere with 5% CO2 and 95% air. Subsequently, the cells grew in fresh medium (RPMI 1640 medium containing 10% FBS) containing different concentrations of doxorubicin, the doxorubicin-GTP conjugate, or the doxorubicin-dGTP conjugate for additional 72 hours. After the 72 hours of incubation, 10 μL of 3-[4,5-dimethylthiazole-2-yl]-2,5-diphenyltetrazolium bromide (MTT; 5 mg/mL) was added to each well and incubated for 4 hours at 37 °C. The supernatant was then removed. 100 μL of acidified isopropanol was added to solubilize the MTT-formazan products for 15 min on an orbital shaker. Absorbance at 590 nm was measured with a microplate reader (BioTek, Synergy). Cell survival rate (r) was calculated with the following equation: 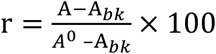, where A_0_ and A represent the absorbance of wells with cells in the absence and presence of a compound, respectively ^49^. A_bk_ is the absorbance of wells without cells. The half maximal inhibitory concentration (IC_50_ value), the concentration of a compound that inhibits 50% cell growth, was calculated according to the cell survival rate.

### Computational Method

The reaction between doxorubicin and GTP was explored theoretically via density functional theory (DFT) calculations of thermodynamic functions for the formation of a van der Waals complex between the two molecules and a chemically bound product through the formation of a new C-N bond between the amino nitrogen in GTP and the carbon of the CH2OH group in doxorubicin with a release of a water molecule. Geometry optimization of the reactants and products was carried out with the B3LYP^50, 51^ functional and 6-31G(d) basis set and vibrational frequencies were computed at the same B3LYP/6-31G(d) level of theory. Next, single-point energies of each molecule were refined using a more accurate long-range corrected ωB97XD^52^ functional and Dunning’s correlation-consistent cc-pVTZ basis set^53^. Finally, solvent effects in an aqueous solution were treated by the self-consistent reaction field (SCRF) method in Truhlar’s SMD implementation^54^. All DFT calculations were performed using the Gaussian 16 quantum chemistry software package^55^. The typical accuracy at this level of theory is within 3-5 kcal/mol.

## AUTHOR INFORMATION

**Corresponding Author**

## Author Contributions

F.L. designed research; G.M., L.S., P.P., K.J.D., and A.M. performed research; F.L., A.M.., F.F.-L., J.H., and A.M. analyzed data; F.L. wrote the paper.

