## Supplementary materials for "Anticancer drug doxorubicin spontaneously reacts with GTP and dGTP"

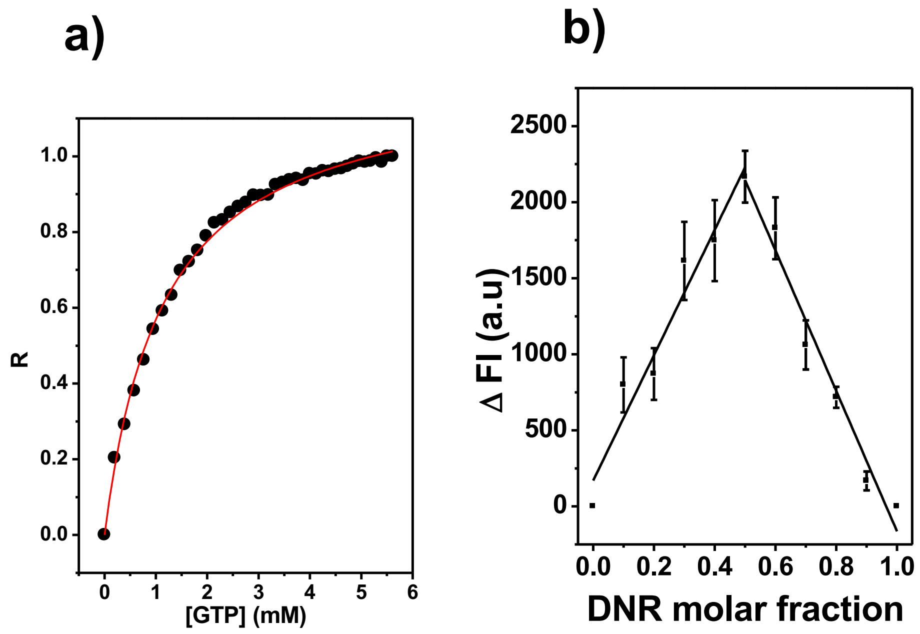


**Figure S1.** Noncovalent binding of doxorubicin to GTP in 10 mM sodium borate buffer pH 8.2. (**a**) The fluorescence titration of doxorubicin to GTP. Data were extracted from Fig. 5D. (**b**) The Job plot for doxorubicin binding to GTP using fluorescence titration experiments.

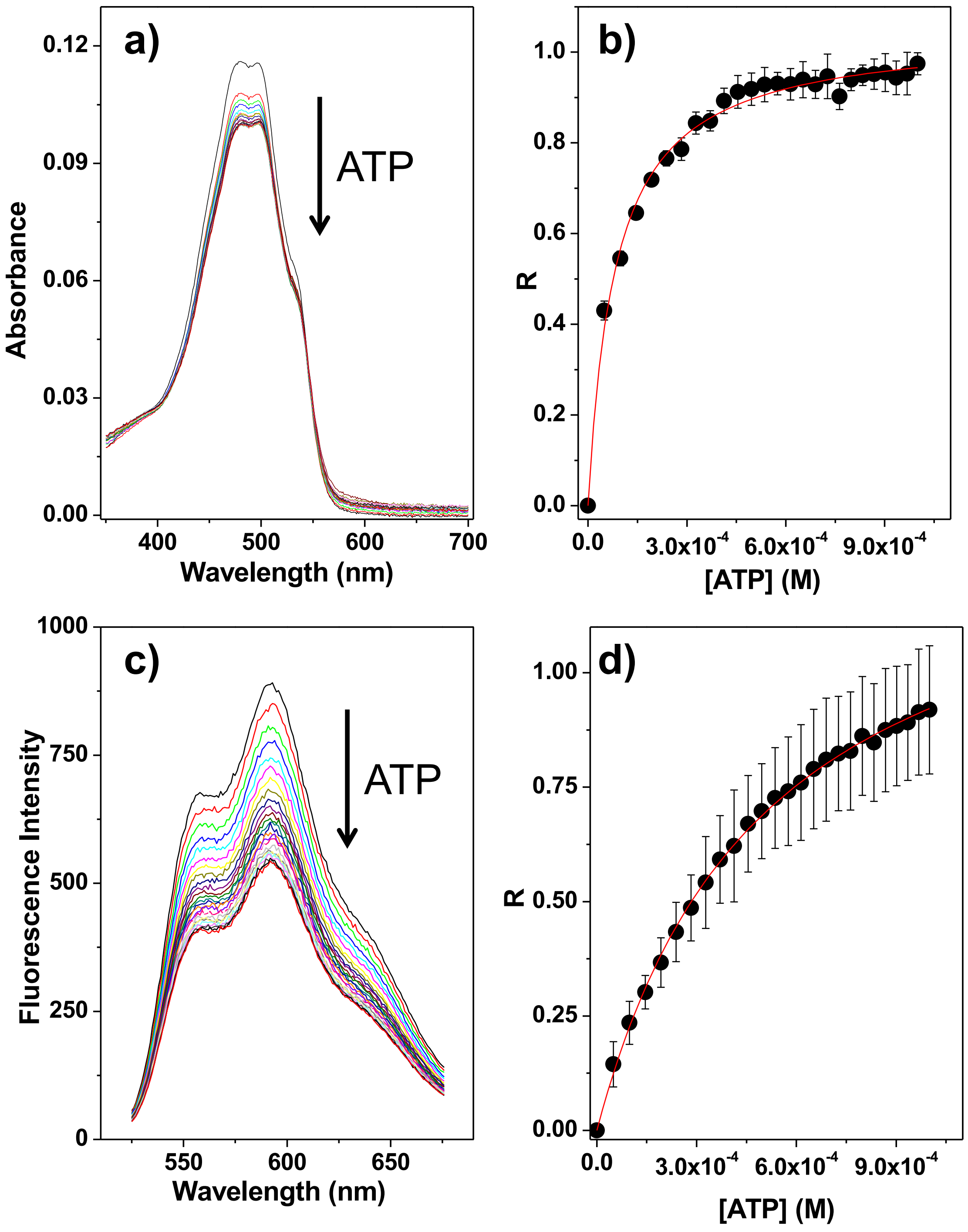


**Figure S2.** Binding of doxorubicin to ATP in 1×BPE buffer. (**a**) Absorbance titration experiments were performed as described under Materials and Methods. (**b**) The binding curve is obtained by nonlinear-least-squares fitting of equation (1). The binding constant of doxorubicin binding to ATP was determined to be $1.2 \pm 0.1\times10^4$ $M^{-1}$. (**c** and **d**) The fluorescence titration of doxorubicin to GTP. Error bars represent standard deviations from three independent experiments.

**Table S1. Binding of doxorubicin to different nucleotides**

| Nucleotide | K ($M^{-1}$) |
|---|---|
| GTP | $3.6 \pm 0.8 \times 10^4$ |
| dGTP | $3.8 \pm 0.8 \times 10^4$ |
| GMP | $2.0 \pm 0.3 \times 10^4$ |
| dGMP | $9.2 \pm 0.4 \times 10^3$ |
| ATP | $1.2 \pm 0.1 \times 10^4$ |
| dATP | $1.4 \pm 0.2 \times 10^4$ |
| dCTP | $1.1 \pm 0.1 \times 10^4$ |
| dTTP | $7.1 \pm 0.7 \times 10^3$ |
| UTP | $4.4 \pm 1.2 \times 10^4$ |

The binding constants of doxorubicin to different nucleotides were determined using absorbance titration experiments as described under Materials and Methods. The data were fitted to a a simple binding model to yield a 1 to 1 binding ratio for all nucleotides.
